# Validation of Dynamic Bayesian Optimization for a Non-Stationary Human-in-the-Loop Optimization Problem

**DOI:** 10.1101/2025.09.08.674907

**Authors:** GilHwan Kim, Fabrizio Sergi

## Abstract

Human-in-the-Loop Optimization (HILO) has demonstrated efficacy in achieving a plethora of assistive or augmentative effects. However, conventional optimizers such as Bayesian optimization (BO) do not account for the non-stationary aspects of the human-robot system, and may thus be limited in the domains of robot-assisted training or rehabilitation.

In this study, we implemented HILO using dynamic Bayesian optimization (DBO) to define the optimal value of a single control parameter to target a desired effect in propulsion mechanics, specifically in the maximum hip extension (HE) angle during stance. Sixteen healthy participants received unilateral hip torque pulses via a hip exoskeleton for about 15 minutes while walking on a treadmill. Exoskeleton torques were determined via HILO using DBO or BO. Unknown to the optimizers, treadmill speed was gradually increased to amplify the non-stationary behavior of the system.

Experimental results revealed that DBO outperformed BO in the later stages of training, leading to improved cost and to final values of torques closer to the true value. The primary difference between optimizers arose from their ability to model the history-dependent relationship between applied torque and output. BO associated later deviations between expected and measured output to process noise, while DBO developed a more accurate predictive model of the input-output relationship that gave proper weighing to past datapoints for making predictions.

## I. INTRODUCTION

Robot-assisted gait training offers the advantage of delivering consistent and repeatable mechanical actions during walking, compared to standard training methods [1], [2]. In such training, objective training effects can be achieved by applying an appropriate level of robotic assistance to the user. Typically, the level of robotic assistance is determined by identifying the user’s input-output relationship or by relying on group-averaged responses. However, estimating individual responses using group-averaged model eliminates inter-individual variability during construction of a response model, which can result in sub-optimal level of assistance for specific participants whose optimal assistance is substantially different from the group average. Alternatively, identifying an accurate input-output relationship model for an individual requires a large number of observations, a number that increases further when the robotic device has large number of control parameters, with a consequent high risk of inducing fatigue. Therefore, efficiently gathering sufficient observations from individuals to construct such input-output relationship model remains a primary challenge in robot-assisted gait training.

To simultaneously account for variability of individual responses and reduce the number of required observations to generate a specific individual response model, Human-in-the-loop Optimization (HILO) has been introduced [3], [4]. By iteratively tuning control parameters during training, HILO enables efficient identification of a individual-specific response model and corresponding optimal input, using different optimization methods that include gradient descent, Bayesian Optimization (BO), and Covariance Matrix Adaptation Evolution Strategy [5], [6], [7], [8].

Previous implementations of HILO in human-robot interaction scenarios targeting walking has primarily targeted the minimization of metabolic cost. While effective for human augmentation purposes, this approach requires relatively longer bouts of walking to establish a reliable estimation of the cost function at each iteration compared to other metrics such as propulsion mechanics or walking speed. Moreover, metabolic cost is not a primary outcome of studies that focus on motor rehabilitation, which may target more directly aspects of biomechanical performance, including walking speed or propulsion mechanics.

Although HILO has the potential to deal with non-stationary system, such as changes in individual response due to neuromotor adaptation or learning, optimizer forms used in prior implementations have relied on the assumption that the human-robot system remains stable over time or across interventions. This assumption disregards the influence of training history on the response of an individual, leading the optimizer to expect similar outcomes when applying identical torque/force assistance regardless of the timing or prior training. Such assumption limits the ability of previous implementations of HILO to account for the changes in individual participant during training.

Dynamic Bayesian optimization (DBO) was introduced in previous studies to support identification of optimal inputs in problems where the cost function could change over time. In DBO, this functionality is achieved by explicitly accounting for the time of inputs and output in the model covariance function [9], [10]. While DBO has been validated in prior studies that aimed to identify the optimal input for time-varying systems in simulations [9], [10], [11], DBO has not been tested in the context of HILO for robot-assisted gait training. Previous studies showed the implementation of DBO in HILO simulations involving neuromotor learning [12] and in a pilot experimental study involving robot-assisted gait training [13], showing performance that was similar to or exceeded that of standard BO. For accurate comparison of controller performance across different implementations and thus HILO runs, it is critical to compare outcomes not only during optimization, but also in separate cross validation sessions. In absence of such validation trials, hypothetical differences in performance measured across optimizers may arise due to the inherently different exploration-exploitation trade-offs pursued by different optimizers, a trade-off that is impossible to control over different optimizer implementations. Therefore, to fully establish the performance of different HILO implementations in non-stationary problems, it is important to periodically compare performance longitudinally in repeated cross-validation iterations, an experimental design that has not been pursued in previous HILO experimental studies.

In this study, we implemented HILO on a hip exoskeleton using DBO to define the optimal value of a single control parameter targeting a desired effect in propulsion mechanics over a single sessions lasting approximately 15 minutes, and compared performance between DBO and a standard BO algorithm. We hypothesized that DBO would be more suitable for identifying optimal solution in a training protocol featuring non-stationary aspects. In a non-stationary problem, the target outcome measure would need to exhibit substantial variability due to neuromotor adaptation and learning, or the participant would have to experience training for a sufficiently long time to result in a significant change in individual response. To establish the feasibility of DBO for such a non-stationary problem in a convenient experimental model, while avoiding fatigue, we decided to amplify the non-stationary aspect of the human-robot system by gradually increasing treadmill speed during training. To compare the performance across optimizers, we incorporated validation iterations in which the optimizer tested its best estimated input value, expected to minimize cost function value. Experimental results showed that the DBO outperformed BO during the later stage of training.

## II. Methods

### A. Dynamic Bayesian Optimization

DBO is a modified version of BO. A detailed description of the method is provided in previous studies [11], [12], but a general overview of the DBO framework is provided in this section for the purposes of following the experimental work presented in this paper. In general, BO uses a Gaussian process (GP) model to estimate outputs corresponding to unobserved input values. Based on this probabilistic estimation model, BO iteratively searches for the input value that minimizes a cost function that is evaluated under measurement noise [14]. The GP model is regenerated iteratively when new observation is provided. The input value (or set of values) to be tested at the next iteration is determined by maximizing the acquisition function. This acquisition function scores candidate input values based on the predicted performance and the related uncertainty, so to balance exploration and exploitation.

The conventional BO framework assumes that the system is stationary; i.e. BO expects similar outputs when inputs are repetitively applied to the system. However, this assumption becomes inaccurate when the system includes a dynamic component, as it is the case when human factors are involved due to neuromotor learning and adaptation [15]. In standard BO, the GP model relies on a stationary covariance function *k*(*u, u*′) between inputs *u* and *u*′, which leads to similar predictions for repetitive inputs. To account for the dynamics of the system, in DBO, the GP model was modified by introducing the time variable *t* into the covariance function, resulting in *k*((*u, t*), (*u*′, *t*′)), where *t* and *t*′ represent the time instances associated with the inputs *u* and *u*′ respectively. This covariance function has been simplified in previous implementations of DBO [10], [16] by assuming that the components of the covariance function are separable, allowing the overall covariance function *k*((*u, t*), (*u*′, *t*′)) to be expressed as a multiplication of a static *k*(*u, u*′) and dynamic component *k*(*t, t*′):

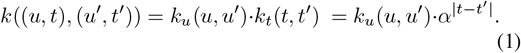

In this work, *k*_*u*_(*u, u*′) was implemented as the automatic relevance determination squared exponential function [17]. Hyperparameter *α* ∈ (0, 1] defines the time scale in the sense that the covariance between measurements collected at times *t* and *t*′ is scaled by a factor *α*^|*t*−*t*′|^ (Fig. 1b).

**Fig. 1.**
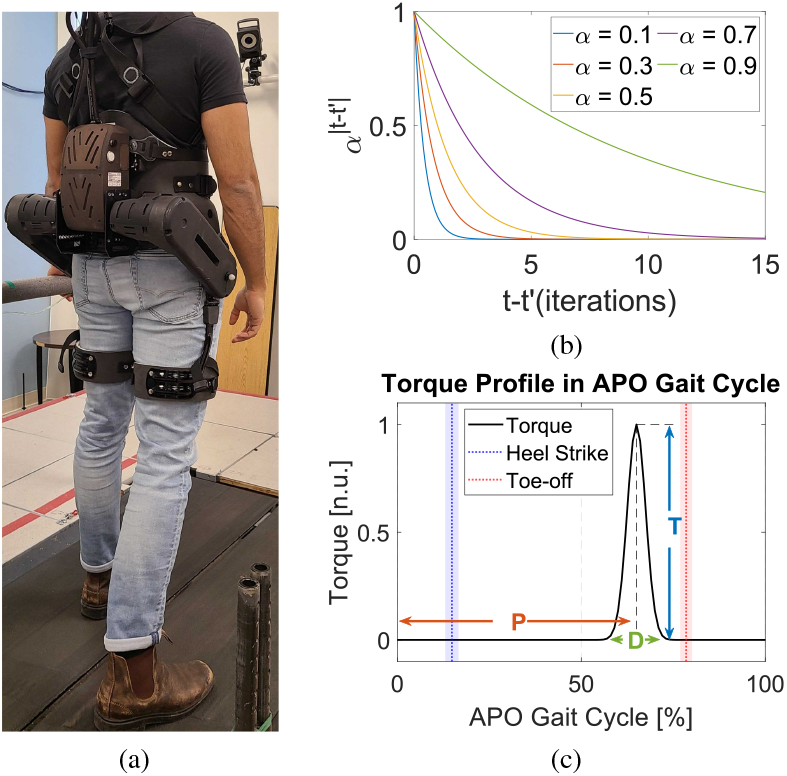
(a) Experimental setup including the active pelvis orthosis (APO), instrumented treadmill, motion capture system, and safety harness. (b) Dynamic component of the covariance function. Each line represents the covariance function with different hyperparameter *α*. (c) Hip torque profile in the APO Gait Cycle domain where 0% is peak of hip flexion. Heel-strike and toe-off timings are shown as distributions, with the mean and standard deviation indicated in blue and red, respectively. The control parameters defining the torque profile (duration: *D*, timing: *P*, and amplitude: *T*) are labeled.

### B. Experimental Methods

#### 1) Equipment

A lower extremity robotic exoskeleton, the Active Pelvis Orthosis (APO), was used to provide torque pulses to participants while walking on the treadmill (Fig. 1a). The APO (IUVO, Pisa, Italy) is a bilateral hip exoskeleton with a weight of 6 kg including battery [18]. An onboard encoder in the APO was used to measure the hip angle with a resolution of 0.015 deg and sampling rate of 100 Hz. Based on the hip joint angle measurements, the APO estimated the participant’s gait phase via adaptive oscillators [19], [20]. These estimates were computed individually for the right and left legs. Using the predicted gait phase and hip kinematics, hip torques were applied in the form of a Gaussian function shaped pulse profile, defined by three control parameters: torque pulse amplitude (*T*), duration (*D*), and peak timing (*P*) (Fig. 1c).

In this experimental study, a single hip torque pulse was applied to the participant’s right hip during each stride. The hip torque pulse was applied near the end of the stance phase, with the duration (*D*) of 18% of the gait cycle (full width at half-maximum: 7.07%), and the timing of peak torque amplitude (*P*) of 65% in the APO gait cycle (0% at peak of hip flexion), resulting in the peak torque being applied at the end of the stance phase. The torque pulse amplitude (*T*) was iteratively updated in real-time via Simulink (MathWorks Inc., Natick, MA, USA). The Simulink controller communicated with the APO via a Serial Peripheral Interface connection. To ensure proper execution of the updated torque profile, the new amplitude command was sent to APO during the swing phase, so that the new torque pulses would be applied in the following stride during the stance phase.

Analog/force plate data from both the right and left side were collected via an instrumented split-belt treadmill (Bertec Corp., Columbus OH, USA) that houses two independent force plates located beneath each treadmill belts. A Simulink-based controller was used to collect signals from the force plates and to send speed commands to the treadmill at 400 Hz.

Leg kinemmatics were were collected in real-time via a 10-camera motion capture system; T40-S (Vicon Motion Systems Ltd, Oxford, UK) with primary goal of tracking the trailing limb angle (TLA) of both legs. Cameras were used to track four retroreflective markers placed on the participant: right and left side of malleoli, and on both side of the rotational axis of APO which was aligned with the participant’s hip joint. Marker locations were recorded at a sampling rate of 100 Hz. For safety, participants were secured using a harness system (Solo-Step Inc., North Sioux City, SD, USA) connected to an overhead railing track during experiment.

#### 2) Study Participants

16 healthy participants (8 females; age: 25.6 ± 2.1 yrs, height: 170.3 ± 7.0 cm, and mass: 73.4 ± 10.3 kg) were recruited for this experiment (protocol no. 1755609-7, approved by our institution’s Institutional Review Board). Each participant conducted two training sessions on the same day, with a rest period of at least fifteen minutes between sessions to prevent fatigue and minimize potential remaining training effects of the first training session. The two sessions differed in the optimization algorithm used (DBO vs. BO), and the order of optimizer used for a participant was randomized and evenly balanced across participants.

#### 3) Pre-training Procedures

The experimental protocol consisted of a pre-training phase followed by two training sessions. During the pre-training phase, retroreflective markers were placed on the participant and APO. The participant donned the APO, aligning the rotational axis of the APO with the participant’s hip axis in sagittal plane. Once the participant confirmed comfort, the device and safety harness was secured. The participant then walked on the treadmill at low speed (0.55 m/s) for approximately 2 minutes with the APO in transparent mode, where the APO sought to minimize the interaction force/torque between the device and user. Treadmill speed was then gradually increased by 0.05 m/s every 5 strides, maintaining a constant speed over such intervals, until reaching 1.2 m/s. The maximum hip extension (HE) angle of the right leg during stance phase from the last 3 strides at each treadmill speed was collected and used to estimate the linear relationship between HE and treadmill speed for each participant. Based on the estimated linear model, treadmill speed required to induce the target change (5 degrees) in participant’s HE was calculated. Immediately after collecting HE data at different treadmill speed, the treadmill speed was set to 1 m/s, and the familiarization session was conducted. During this session, participant experienced a series of torque pulses (extension: 1, 3, 5, 7, 9 Nm, flexion: 1, 3, 5 Nm) applied in ascending order for ten strides each (duration *D* = 18% and timing *P* = 65%, as tested during optimization). Following the pre-training phase, participants were given a rest period approximately 5 minutes.

#### 4) Treadmill Speed Increment During Training

Given their formulation, DBO and BO may perform differently if the system response is sufficiently non-stationary [10], [11], [12]. In practical training applications, such difference may emerge when the target outcome is highly non-stationary or when participants are exposed to training over a sufficient number of iterations, possibly over multiple repeated sessions. However, testing a new algorithm on a multi-session study conducted in natural walking conditions would be an inefficient experimental design that would be overly burdening for participants. As an initial step towards validating the feasibility of a optimizer algorithm in HILO that can account for the non-stationary properties of the human response to training, we opted to intentionally induce a controlled amount of non-stationary behavior in the measured response.

Because most metrics of walking mechanics are speed-dependent, we opted to gradually increase treadmill speed during training, a condition that was obviously completely unknown to the optimizer. In this condition, the optimizer would be required to realize that the optimal input in the initial stages of the experiment would be substantially different from the one that would be required in the late stages of the experiment. Moreover, because the treadmill speed conditions were consistent across optimizer/session for a given participant (BO vs. DBO), this experimental design allowed us to directly compare outcomes between optimizers during training.

The overall speed range throughout for whole training was determined based on the outcomes of the pre-training session. The maximum (final) speed was fixed at 1.2 m/s, and each participant’s initial speed (low speed) was calculated by subtracting the individual speed range from 1.2 m/s. Based on the observation that maximum HE increases with walking speed, the speed range was derived from a linear model based on pre-training session data to produce equivalent changes in the target outcome (5 degrees). If the calculated low speed fell below 0.55 m/s, it was set to 0.55 m/s. The speed range was defined such that the ideal torque input was expected to decrease progressively over the course of training, ultimately reaching zero torque amplitude by the final iteration (80th iteration).

Treadmill speed was initially set to the low speed for baseline and the first iteration. Speed was then increased by a fixed increment at each iteration to reach 1.2 m/s by the 80th iteration. Treadmill speed remained constant within iteration and was updated right after cost function evaluation. For example, at the i-th iteration, the cost function value was calculated from observations of the 1st to 5th strides. Immediately following 5th stride, the treadmill speed was updated.

#### 5) Training Procedures

The training session consisted of a baseline phase followed by a torque intervention phase. During the baseline phase, participants walked at low speed in transparent mode of APO, without any torque pulses, for 110 strides. The torque intervention phase started right after. During this phase, torque pulses were applied to the right hip joint at the end of the stance phase.

The torque intervention session consisted of 80 continuous optimizer iterations, where each iteration consisted of either 6 or 7 strides, depending on the time it took for the optimizer to identify the input for the next iteration. Within each iteration, the APO applied the same torque pulse continuously from the first stride until the optimizer updated the control parameters for the next iteration. Based on the participant response, the optimizer updated the control parameter *T*. HE was measured over the first 5 strides of each iteration, and the average value of these 5 measurements was used to calculate the cost function value *c*(*i*), which was sent to the optimizer. *c*(*i*) was defined as the absolute difference between the desired HE and the average HE measured at that iteration. After completing the first training session, the participant removed all equipment, including exoskeleton and harness, and was asked to rest for at least 15 minutes before starting the second training session with the different optimizer.

#### 6) Optimizer Setup

The objective of both optimization methods (DBO and BO) used in each training session was to increase maximum HE of the right leg during stance phase by 5 degrees relative to baseline. The baseline HE (*HE*_*BL*_) was calculated by averaging maximum HE during stance phase over the last 20 strides of the baseline session. At each iteration *i*, HE (*HE*(*i*)) was calculated as average of the first 5 consecutive strides within that iteration. Accordingly, the cost function was defined as absolute difference between desired HE target (*HE*_*BL*_+5°) and measured HE: *c*(*i*) = | *HE*_*BL*_+5° − *HE*(*i*) |. During each iteration, the participant experienced the same torque input for each stride until the optimizer determined the new control input to test. This update took an additional 1-2 strides in this experiment.

This experiment featured a single-parameter optimization method, targeting the parameter of torque pulse amplitude *T*. The range of *T* was from −5 Nm (maximum flexion) to 9 Nm (maximum extension). The input values for the first three iterations were predefined as 5 Nm, 7 Nm, and 3 Nm, respectively. The acquisition function used in this study was expected improvement, selected based on its highest performance in previous simulation results [21]. The exploration-exploitation ratio (e-ratio) was set to 0.1 (on a scale from 0 to 1), favoring exploitation than exploring uncertainty. This optimizer value was set based on pilot testing, which showed no significant difference between different e-ratio of 0.1, 0.2, 0.4, 0.6, and 0.8 when using the same optimizer (DBO or BO). Pilot testing indicated that optimization runs with an e-ratio of 0.1 were less overly variable and more intuitive for the user.

A total of 80 iterations were performed in each training session, with validation iterations conducted every 10th iterations. For instance, at the 60th iteration, the optimizer used input and output data from iteration 1 to 59 to estimate the best input value to induce desired change from participant’s response, which was then tested in 60th iteration. This procedure allowed to validate the accuracy with which the optimizer was able to form a model of the response between input and output at the estimated optimal input, and compare between optimizers without confounds related to different instances of exploration pursued by different optimizers or optimization runs. Based on the GP model, the input value with the minium upper confidence bound of the cost function value was selected as the estimated best input value.

#### 7) Statistical Analysis

Individual participant responses at each validation iteration during training were collected and analyzed. Since treadmill speed was matched across both optimizers for all participants, paired comparisons of outcome measure at each validation iteration were used to assess the performance of the two optimizers. Outcomes subjected to statistical analysis included the following primary outcomes, which were expected to directly be influenced/modulated by the optimizer: change in HE relative to baseline, cost function value, and applied torque. Moreover, secondary outcomes such as cadence, normalized peak anterior ground reaction force (nAGRF), normalized propulsive impulse (nPI) integrated over the positive portion of the AGRF curve, the maximum trailing limb angle of the right leg during stance (max. TLA), were also analyzed in a similar fashion to assess generalization to other components of walking not directly targeted by the optimizer. Finally, the prediction error of the best-estimated response at the cross-validation iterations was also calculated to assess the accuracy of the model developed by BO and DBO longitudinally across an experiment.

To assess the need of parametric testing, data distributions for each outcome across the sixteen participants at each validation iteration were evaluated for normality using the Shapiro–Wilk test. Pairwise t-tests were conducted to compare outcome differences between optimizers at each validation iteration. In addition, two two-way repeated-measures ANOVAs were performed to examine the effects of optimizer type (DBO vs. BO) and iteration number (20th vs. 80th) on cost function values and applied torque, with the 20th and 80th iterations representing early and late training phases, respectively. A repeated-measures design was chosen to account for within-subject variability, as the same participants experienced both optimizers across multiple iterations. When significant main effects or interactions were identified, post-hoc pairwise t-tests were performed to compare each combination of two factors:optimizer and iteration number.

## III. Results

### A. Performance difference between DBO and BO

Group averaged cost function values at each iteration are shown in Fig. 2. Results from each optimizer (DBO and BO) were subject to paired comparisons at the validation iterations, where each optimizer tested the best estimated input value. As shown in Fig. 2, the performance of the two optimizers was indistinguishable until the final iterations (specifically, iterations 70 and 80). At the 70th iteration, DBO resulted in a cost function value of 1.5 ± 1.0° (mean ± std), while the cost for BO was 3.0 ± 1.2°, with a significant difference between DBO and BO (1.5 ± 1.5°, *p* = 0.002). The difference between DBO and BO remained significant also at the final validation iteration (1.8 ± 1.4°, *p* = 0.002), with DBO resulting in a cost function value of 1.8 ± 1.4°, and BO resulting in a cost function value of 3.5 ± 1.3°.

**Fig. 2.**
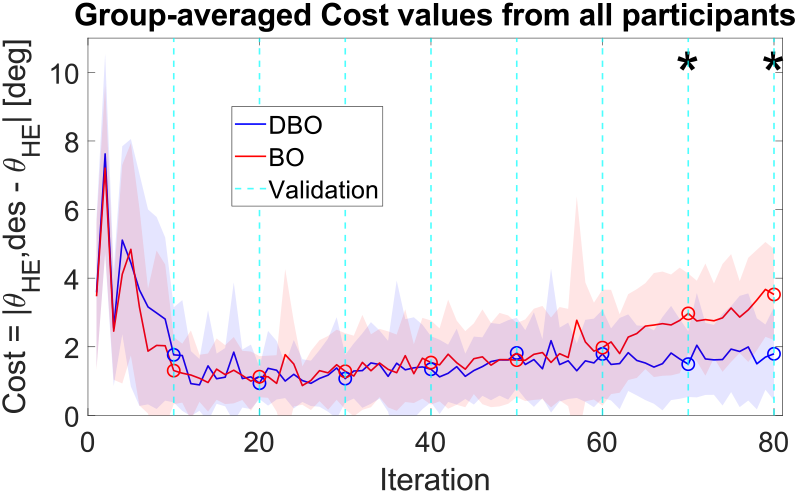
Group level cost function values during the experiment. The solid line indicates the mean cost function value for 16 participants at each iteration, and the shaded area extends by one standard deviation. The cyan dashed line indicates validation iterations where the estimated best input was tested, corresponding cost function values are shown with a circle marker. An asterisk represents significant difference between DBO and BO (p *<* 0.05).

Group averaged torque amplitude at each iteration is shown in Fig. 3. In BO, the mean torque amplitude remained relatively constant from the 10th iteration to the end of experiment. In contrast, DBO showed a progressive decrease in average torque amplitude over iteration, a trend closer to the one expected due to the fact that speed progressively increased during training. Significant differences in torque amplitude between DBO and BO were observed at the 70th (1.0 ± 1.5 Nm, *p* = 0.019) and 80th (1.4 ± 1.5 Nm, *p* = 0.001) validation iteration. At the 70th iteration, DBO applied a significantly lower torque amplitude (1.6 ± 1.6 Nm) compared to BO (2.6 ± 1.0 Nm). The difference remained significant at the 80th iteration (DBO: 1.2 ± 1.3 Nm, BO: 2.6 ± 1.0 Nm).

**Fig. 3.**
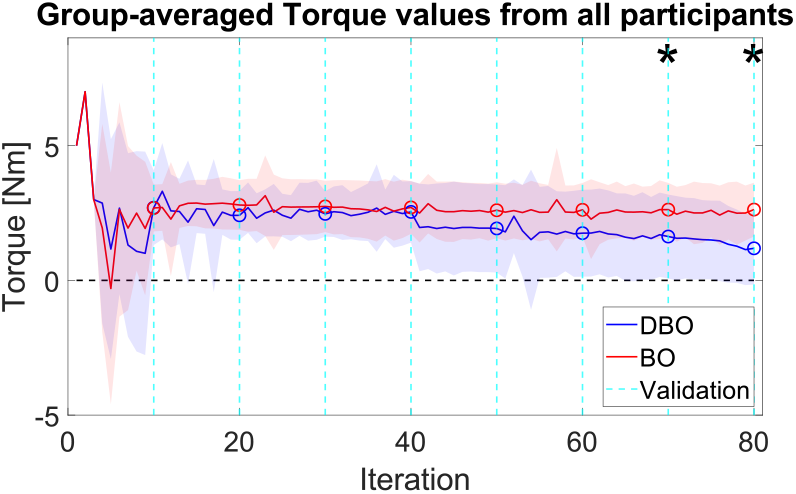
Group level torque amplitude at each iteration. The black dashed line indicates the zero value, which is the expected value that the optimizer should reach at the end of the experiment. Torque amplitude is positive in the direction of hip extension.

The two-way ANOVAs were conducted to analyze longitudinal changes in cost function value and torque. The ANOVA for cost highlighted that both optimizer type (*p <* 0.0001) and iteration number (*p <* 0.0001) significantly affected cost function value, as did the interaction between optimizer type and iteration number (*p* = 0.004). As shown in Fig. 4, post-hoc tests indicated that cost increased slightly between iterations for both DBO and BO (DBO: 0.9 ± 1.5°, BO: 2.4 ± 1.2°, but the increase in cost between iterations was greater for BO than it was for DBO (1.5 ± 1.7°, *p* = 0.005). The ANOVA for torque highlighted a significant effect of both factors (optimizer type: *p* = 0.001; iteration number: *p* = 0.011), but no significant interaction effect. Similarly as for cost, the amount of applied torque decreased slightly between iterations for both DBO and BO (DBO: −1.2 ± 1.6 Nm; BO:−0.2 ± 0.3 Nm), and remained significantly different from zero at iteration 80 (DBO: 1.2 ± 1.3 Nm, *p* = 0.003; BO: 2.6 ± 1.0 Nm, *p <* 0.0001), however the decrease in torque was significantly greater in magnitude for DBO compared to BO (1.1 ± 1.7 Nm, *p* = 0.02). All individual responses for cost function value and applied torque from the experiment are shown in Fig. 6.

**Fig. 4.**
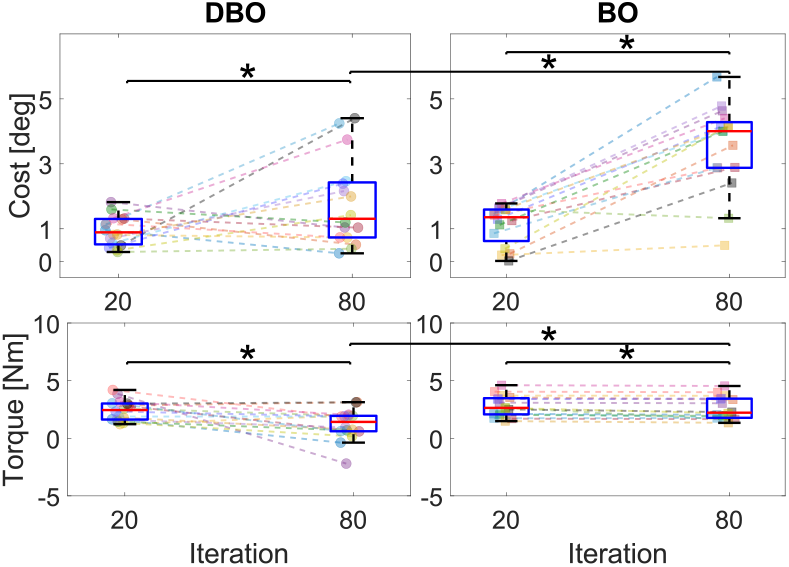
Paired comparisons broken down by iteration number and optimizer type. Colored dots indicate each participant’s data. An asterisk represents a significant difference based on paired comparisons.

Changes in group-averaged maximum HE at each iteration relative to baseline are shown in Fig. 5. Consistent with the cost function results (Fig. 2), DBO produced significantly different outcomes compared to BO at iteration 70 (1.6 ± 1.8°, *p* = 0.004) and 80 (1.9 ± 1.9°, *p* = 0.001). DBO maintained HE values close to the desired increase relative to baseline (70th iteration: 6.3 ± 1.2°, 80th iteration: 6.7 ± 1.5°). In contrast, BO resulted in a larger deviation from the target 5° increase, with observed values of 7.9 *±* 1.4° at the 70th iteration and 8.5 *±* 1.3° at the 80th iteration.

**Fig. 5.**
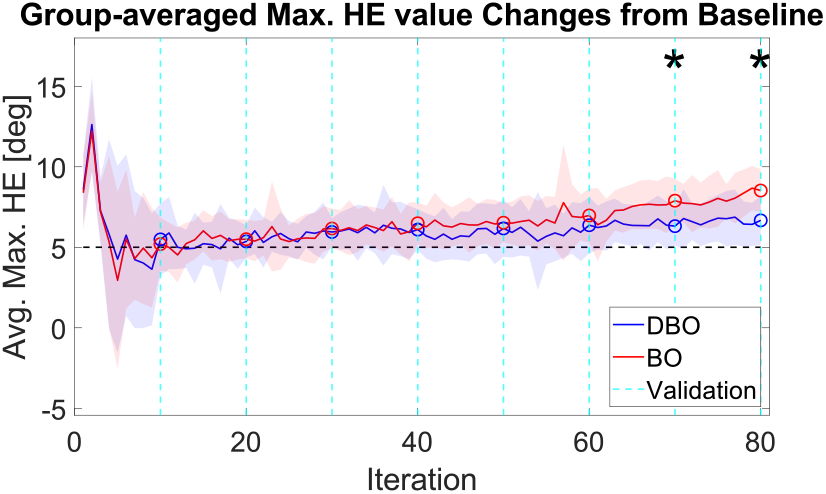
Group level maximum HE change relative to baseline at each iteration. The optimizers have a target of a 5-deg increase in maximum HE relative to baseline (black dashed line).

**Fig. 6.**
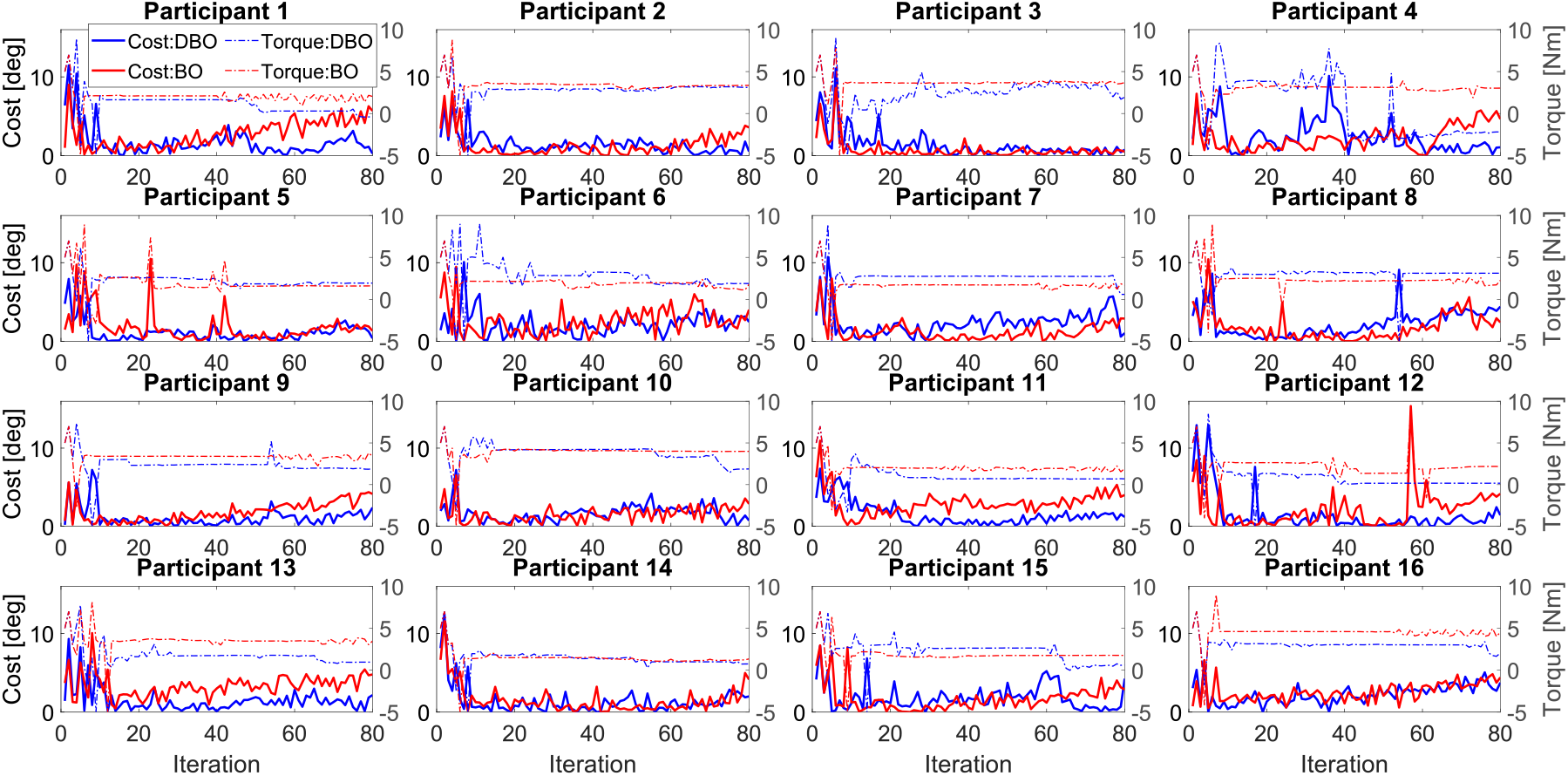
Individual participant responses. The solid line shows the cost function value at each iteration, and dashed line indicates the corresponding torque amplitude applied to the participant.

### B. Training-induced changes in secondary outcomes of propulsion mechanics

Group averaged results for secondary outcomes quantifying propulsion mechanics of the leg exposed to exoskeleton torque are reported in Fig. 7. As expected, all of these outcomes increase as the imposed treadmill speed increases. Unlike the primary metrics reported above, there was no significant difference between the results from DBO and BO.

**Fig. 7.**
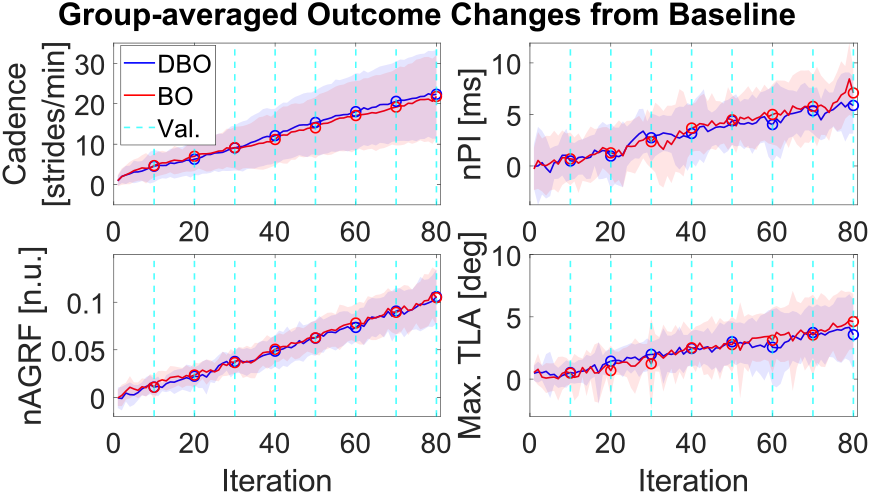
Group level change in secondary outcomes at each iteration relative to baseline. The solid line indicates the mean of all participant’s average outcome change at each iteration (n = 16 for cadence, nPI, and nAGRF, and n=13 for max. TLA), and the shaded area represents one standard deviation.

## IV. Discussion

The main goal of this paper was to validate DBO in a non-stationary HILO problem in the context of robot-assisted training of propulsion mechanics. An experiment involving a robotic hip exoskeleton and sixteen healthy participants was conducted to compare the outcomes of DBO with those of a standard BO algorithm. Results of the experiment indicate that DBO exhibited improved performance relative to BO, primarily because it was more capable than BO of accounting for non-stationary aspects in the human-robot system. This is clearly demonstrated in Fig. 2 and 5, where the results from DBO show smaller values of cost in the later stages of the walking experiment, despite the imposed increase in treadmill speed. In contrast, the results from BO exhibit a form of drift of the cost function value as iterations progress. Statistical analysis revealed a significance difference between DBO and BO results starting from the later stages of training (iteration 70th), which persisted through the final validation iteration.

The non-stationary aspect of human-robot system in this experiment was amplified by gradually increasing treadmill speed, with adjustments individualized based on each participant’s pre-training responses. Since participants were expected to require progressively less torque as training proceeded, ideally reaching zero torque at the end of training, the desired torque amplitude at each iteration was expected to decrease during training. As shown in Fig. 3, group average torque amplitudes from DBO and BO diverged significantly at the 70 and 80th validation iterations. Both optimizer achieved the target HE change early in the training (around iteration 10 and 20, Fig. 5). This early achievement resulted in BO applying similar torque amplitude longitudinally across training (20th iteration: 2.8 ± 0.9 Nm, 80th iteration: 2.6 ± 1.0 Nm). In contrast, DBO escaped from keeping such a constant torque amplitude pattern around the 40th iteration and resulted in a larger overall decrease in torque amplitude during training compared to BO (20th iteration: 2.4 ± 0.9 Nm, 80th iteration: 1.2 ± 1.3 Nm). As shown in Fig. 4, both DBO and BO showed a significant decrease in torque amplitude from the early (20th iteration) to late stage of training (80th iteration), but the reductions were insufficient to reach zero torque at the last validation iteration. As visible in Fig. 5, the applied torques resulted in the outcome measure exceeding the target, implying that both optimizers consistently estimated the optimal input to be greater than the true value. This phenomenon is likely due to the fact that both optimizers have some unavoidable delay in estimating the GP model accurate at the *current* iteration. Given the significant differences between DBO and BO, such delay was considerably smaller in DBO compared to BO. However, the observation that both optimizers applied a torque that was significantly greater than zero (expected value based on the pre-training session) at iteration 80th indicates that such a delay was non-zero for both optimizers.

The performance difference between DBO and BO reflects their differing ability to account for the relevance of past observations to current conditions, which is closely tied to the model used by each optimizer to define an optimal input at each iteration during training. An example describing the training history for a selected participant is shown in Fig. 8. Initially (e.g., iterations 10-20), both optimizers applied small torque amplitude required to achieve the target increase in HE. As training continued, because the treadmill speed increased, the amount of torque necessary to achieve the same outcome should have decreased over iteration, leading to DBO and BO observing outcomes that differed from expected values. While BO interpreted those differences to arise from the unstructured variability of the human-robot system (process noise + measurement noise), leading to an increase in the confidence interval of the estimated GP model at later iterations, DBO was able to correctly assign the source for such a mismatch to the fact that there is a time-dependent co-variance between measurements collected at different time points. As such, DBO was able to correctly down-weigh the observations collected in the early phases of the experiment, in favor of more recent observations that led to better model predictions, resulting in the application of torque inputs that were closer to the expected value of zero, leading to a smaller cost compared to BO. While this is intuitively visible in the single-subject plot (Fig. 8), the prediction errors obtained via DBO and BO are significantly different also at a group level (Fig. 9).

**Fig. 8.**
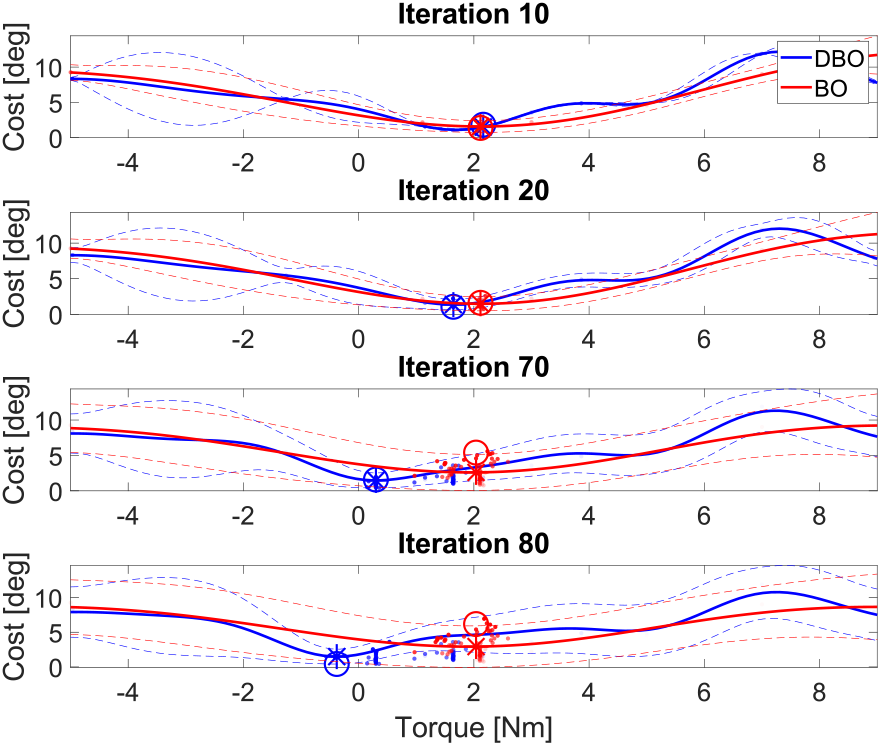
Gaussian process models from participant 1 during training. Models were constructed using the response history before the validation sessions at iterations 10, 20, 70, and 80. The solid line shows the estimated mean cost function across torque values, with dashed lines indicating the 5th to 95th percentile confidence interval. Circles and asterisks denote the estimated mean and actual response at each validation session, respectively. Dots represent past responses, with greater transparency indicating earlier iterations.

**Fig. 9.**
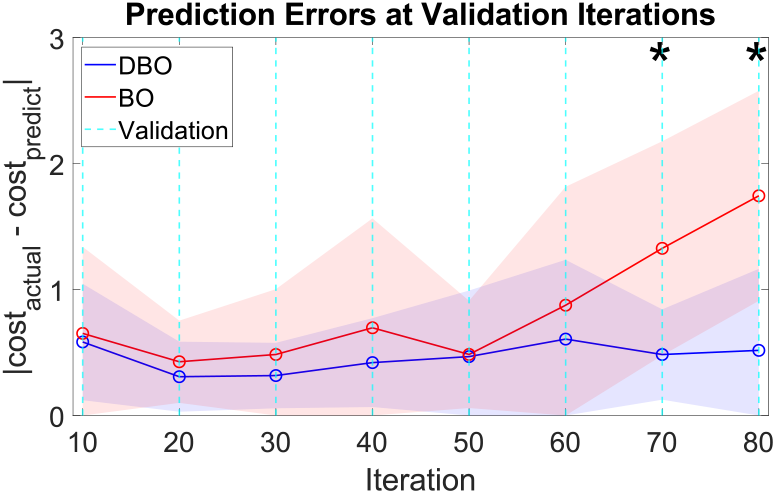
Group-averaged prediction error at each validation iteration. Prediction error was calculated based on the absolute difference between estimated cost value from optimizer and measured cost value. Blue and red markers indicate the mean from DBO and BO respectively, with the shaded area indicating one standard deviation.

## V. Conclusion

This study validates the use of DBO in a non-stationary HILO problem in the context of robot-assisted training of propulsion mechanics. Performance differences between DBO and BO were evaluated using validation iterations, where the estimated best solutions from each optimizer were tested. The results showed that DBO outperformed BO in the later stage of training.

There are some limitations to this study. In real-world application for training propulsion mechanics, the dynamic changes observable within a similar time-span (10-15 minutes of robot-assisted gait training) may be less pronounced than those observed in this study, where a non-stationary component was intentionally introduced by increasing treadmill speed. Consequently, the performance difference between optimizers may be smaller in other settings. However, it is worth noting that some domains of application of human-robot interaction involve sessions lasting hours and include the repetition of multiple sessions [22], [23], making this method potentially a good candidate for extension to scenarios of long-term human-robot interaction in the context of assisted gait training. Additionally, the cost function used in this study, defined as the absolute difference between the desired change and current change in HE at each iteration, is not expected to be a smooth function since it exhibits a discontinuity of the derivative at the optimal input. This characteristic limits the optimizer performance to precisely model the cost function relative to torque using GP modeling.

In future work, the observation from this study will be expanded to include larger number of control parameters targeting more biomechanically relevant outcomes such as walking speed.

